# WTools: a MATLAB-based toolbox for time-frequency analysis

**DOI:** 10.1101/2024.10.31.621249

**Authors:** Ambra Ferrari, Luca Filippin, Marco Buiatti, Eugenio Parise

## Abstract

Electroencephalography (EEG) is an established method for investigating neurocognitive functions during human development. In cognitive neuroscience, time-frequency analysis of the EEG is a widely used analytical approach. This paper introduces WTools, a new MATLAB-based toolbox capable of performing time-frequency analysis of EEG signals using complex wavelet transformation. WTools features an intuitive GUI that guides users through the analysis steps, focusing on essential parameters. Being free and open-source, it can be integrated and expanded with new features, making it a handy tool that is growing its popularity in developmental cognitive neuroscience. Here, we provide a detailed description of the WTools algorithm for wavelet transformation and we compare it with state-of-the-art methods implemented in EEGLAB. Alongside the official tool release, we also offer a comprehensive illustrated tutorial to enhance accessibility and promote the transparent and reproducible usage of WTools by the scientific community.

**Highlights:**

- WTools is a new MATLAB-based toolbox for time-frequency analysis of the EEG signal
- It uses wavelet transformation and the algorithm is fully described
- It is designed to be very user friendly through its simple GUI
- It is open source and flexible as the code is freely available
- We provide a step-by-step tutorial and an example dataset to learn the toolbox

## 1 Introduction

Electroencephalography (EEG) is an established and practical tool for studying brain function and dysfunction across the lifespan. Over the past decades, time-frequency analysis of the EEG has become increasingly popular in cognitive neuroscience. Often implemented via wavelet transformation (Cohen, 2014; Csibra et al., 2000; Jensen and Tesche, 2002), time-frequency analysis attempts to unfold the EEG signal into the frequency domain. Different from pure Fourier transformation, it includes some temporal support, allowing us to see the frequencies in the EEG signal spread over time relative to a time-locked stimulation. Consequently, time-frequency analysis can finely characterize the temporal dynamics embedded in EEG oscillations in terms of frequency, power, and phase (Cohen, 2014). This analysis provides invaluable information on a variety of cognitive processes such as attention (Jensen and Mazaheri, 2010; Klimesch, 2012), learning and prediction of upcoming information (Arnal and Giraud, 2012; Engel et al., 2001), memory (Griffiths and Jensen, 2023; Hanslmayr et al., 2016), language (Drijvers and Mazzini, 2023; Poeppel, 2014; Wang et al., 2012) and motor activation (Brittain and Brown, 2014; Muthukumaraswamy et al., 2004), pointing to the centrality of cortical rhythms in cognition (Buzsáki and Draguhn, 2004; Wang, 2010).

Over the past two decades, time-frequency analysis of the electroencephalogram (EEG) has become increasingly popular in the field of developmental cognitive neuroscience (Begus et al., 2016; Begus and Bonawitz, 2020; Csibra et al., 2000; Kaufman et al., 2005; Maguire and Abel, 2013; Meyer et al., 2020; Southgate et al., 2010; Xie et al., 2018). Yet, most studies in this field currently rely on ERPs and Fourier-based power, possibly because time-frequency analyses can be both computationally intensive and analytically complex (Morales and Bowers, 2022). Different from event-related potential (ERP) analysis, time-frequency analysis can be done with many different mathematical approaches (Fast Fourier Transformation, Hilbert transformation, wavelet transformation) and requires the manipulation of several parameters. Different approaches and small changes in the parameters can lead to very different results.

There is a large number of free software (e.g. EEGLAB: Delorme and Makeig, 2004; MNE-Python: Gramfort et al., 2013; ERPWAVELAB: Mørup et al., 2007; FieldTrip: Oostenveld et al., 2011) and commercial software (NetStation, Brain Vision Analyzer) to perform time-frequency analysis of the EEG. However, commercial software does not disclose the exact algorithm for the signal processing. As a result, it is virtually impossible to understand what the software does exactly, undermining flexibility, transparency and reproducibility. Free software on the other hand can be difficult to use because of the lack of a graphic user interface (GUI), or because they require to manually set several parameters. By developing in complexity and integrating an increasing number of functionalities, these softwares are difficult to use for non-experienced researchers and students.

This paper presents WTools, a new MATLAB-based toolbox for time-frequency analysis of the EEG signal. WTools is designed to be very user-friendly through its simple GUI; it is easy to learn and use because it keeps the number of parameters to manipulate at a minimum; it is open source and flexible as the code is freely available. WTools uses wavelet transformation and the algorithm is fully described. It works in combination with EEGLAB, exploiting its functionalities to import several different EEG data formats, as well as some of the EEGLAB plotting functionalities. Its data structure is derived from ERPWAVELAB (Mørup et al., 2007) and therefore WTools files are fully compatible with ERPWAVELAB. It handles multi-channel time-frequency analysis and multi-subject projects. It computes within-subject time-frequency differences among conditions and plots the results with time-frequency plots, as well as 2D and 3D scalp maps. It is possible to integrate and extend the toolbox with personalized configurations for developmental research (i.e. newborn and infant scalp maps). Finally, WTools includes a function to export numerical datasets into tabulated text files that can be easily analysed with any statistical software. For these reasons combined, an increasing number of studies in the field of developmental cognitive neuroscience have adopted beta versions of WTools before its official release (de Klerk et al., 2016; Kolesnik et al., 2019; Ortiz-Barajas et al., 2023; Piccardi et al., 2020; Pomiechowska and Csibra, 2017; Quadrelli et al., 2021, 2019; Southgate and Vernetti, 2014; Yin et al., 2020). With the present work, we aim to formalize the toolbox, publicly release it together with a clear, step-by-step user-friendly tutorial and therefore promote its usage from the wider developmental cognitive neuroscience community and beyond.

To provide a comprehensive overview of WTools, the paper is structured as follows. First, we describe how WTools implements wavelet transformation and how the signal is processed. Then, we illustrate the main visualization functionalities of WTools. Finally, we compare its time-frequency functionality with state-of-the-art methods implemented in EEGLAB, an established popular free software (Delorme and Makeig, 2004). WTools scripts are made freely available for the users (under GPL-3.0 license) on GitHub (github.com/cogdevtools/WTools). A user-friendly step-by-step tutorial of the entire pipeline for the use of WTools is provided on GitHub, along with an anonymized example dataset (see section “WTools availability”).

## 2 Materials and Methods

### 2.1 WTools description

The processing pipeline (Fig. 1) consists of a sequence of steps, each controlled through a GUI and executed by an independent code module. The paper focuses mainly on the core analytical section of the toolbox. Accordingly, in the following sections we describe the wavelet transformation implemented in WTools. The description is kept at the minimum possible complexity to provide a simple and clear explanation of how WTools transforms the EEG signal into the time-frequency domain.

**Fig. 1.**
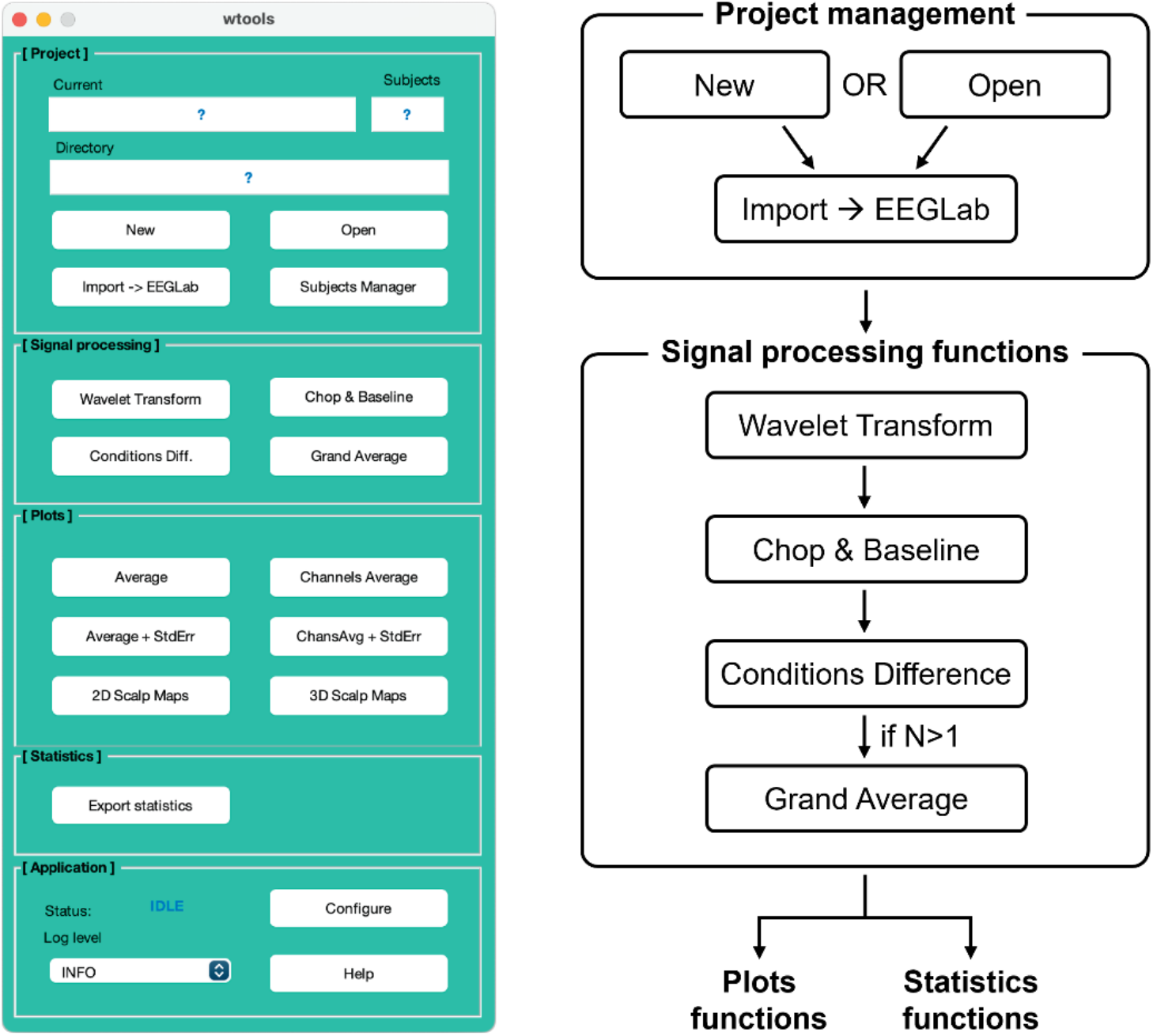
WTools GUI (**A**) and schematic representation of the processing pipeline (**B**). The toolbox is divided into five main sections: Project management (initialize project and import data); Signal processing (run time-frequency analysis); Plots (create time-frequency plots, 2D and 3D scalp maps); Statistics (export numeric values for statistical analyses); Application (configure and monitor the toolbox’s functionalities). The paper focuses mainly on the core analytical section of the toolbox (i.e. Signal processing); for further details on the remaining sections, please consult the step-by-step tutorial accessible from the WTools GitHub project website (see section “WTools availability”).

#### 2.1.1 Morlet complex wavelets

A wavelet (small wave) is a mathematical function of time (*t*) and frequency (*f*), with the following two features. First, its amplitude approaches zero at positive and negative infinity and it differs from zero only around the origin of the cartesian axes. That is, differently from other mathematical functions such as sinusoidal or infinite waves, a wavelet has a defined temporal length. Second, the sum of the area of a wavelet above the x axis is equal to the sum of the area of the wavelet below the x axis, hence the total sum of the area of a wavelet is equal to zero. Therefore, differently from a filter, a wavelet will not change the power spectrum of a signal. WTools computes Morlet complex wavelets at a 1 Hz sampling frequency according to:

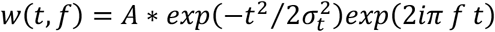

where 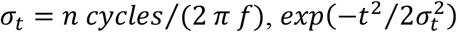 is a Gaussian bell curve, *exp*(2*iπ f t*) is a complex sinusoid. Wavelets are by default normalized so that their total energy is 1, the normalization factor *A*being equal to 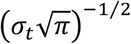 (Tallon-Baudry et al., 1996). This is common practice in time-frequency analysis (e.g. in EEGLAB), which we strongly recommend as it allows direct quantitative comparisons of frequency content across frequency values. Nevertheless, we still provide the option to omit wavelet normalization for back-compatibility with previous (beta) versions of WTools. As a sanity check, there is a perfect match between the wavelets specified by WTools and EEGLAB once wavelet normalization is applied to the WTools wavelet (Fig. 2).

**Fig. 2.**
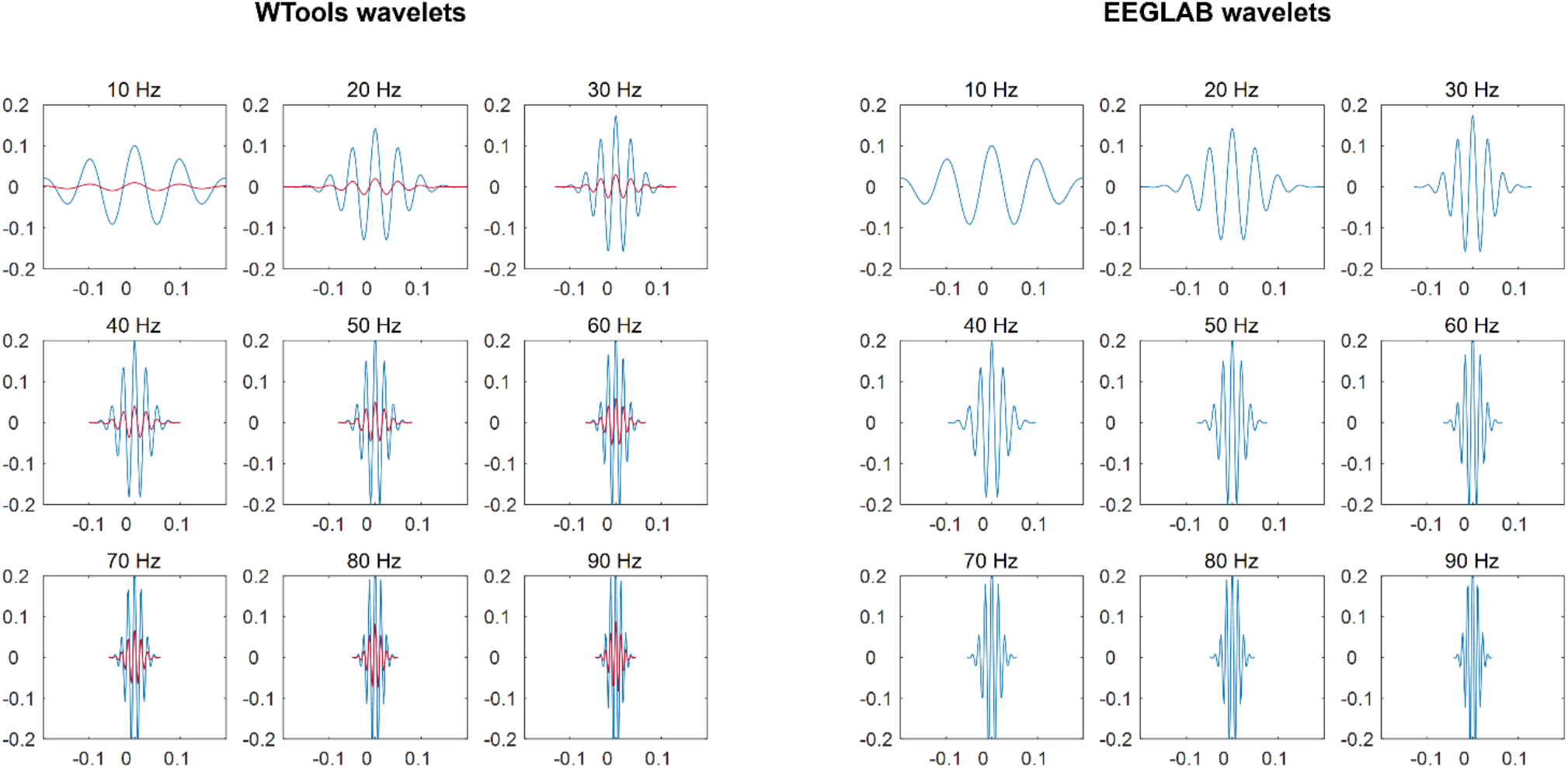
Comparison of complex Morlet wavelets between WTools (on the left) and EEGLAB (on the right) for the frequency range used in the present work (10-90 Hz; see section “Data analysis”). Please note that the standard WTools pipeline computes wavelets with 1 Hz sampling frequency; here we plot wavelets with 10 Hz sampling frequency for visualization purposes only. The red plots show non-normalized wavelets for WTools. The blue plots show normalized wavelets for WTools and EEGLAB, with a perfect correspondence between the two. X axis: Time (s); Y axis: Wavelet amplitude (a.u.).

The last term of the equation can be written as a sum of *sin* and *cos* waves:

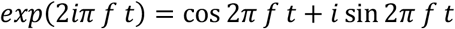

In practice, a wavelet is a complex sinusoidal wave enveloped into a Gaussian bell curve. Because a wavelet is a function depending on a frequency *f*, at each *f* the corresponding wavelet works as a magnifier that can reveal how much that frequency is present into the EEG signal. At the same time, because a wavelet has a defined temporal length, it can preserve some time information after the transformation of the signal.

Wavelets are wide at lower frequencies, that is the temporal information is relatively imprecise but the frequency information is preserved. Wavelets become progressively narrower at higher frequencies, that is the time information is more accurate but the frequency information is less precise. Time and frequency information respond to Heisenberg’s indetermination principle: the more accurately we know one of them, the less accurately we know the other. Wavelet transformation attempts to find the best possible compromise to squeeze as much information as possible in terms of both time and frequency, depending on the frequency of interest.

#### 2.1.2 Time-frequency transformation

After computing all the wavelets for each frequency in a range of interest, WTools uses continuous wavelet transformation. By convolving the signal *s(t)* (cleared of its DC component) with the wavelet *w(t, f)* and taking the modulus of the resulting complex coefficients, WTools obtains the coefficients (the mean square root) of the wavelet transformation at a specific frequency *f*. The resulting vector represents how well the signal fits the wavelet at a the frequency *f*, that is how much of the frequency *f* is in that signal (Herrmann et al., 2005). WTools performs the convolution for each experimental condition and channel, of each EEG segment and subject, for all frequencies of interest. Transformed segments are then averaged across trials:

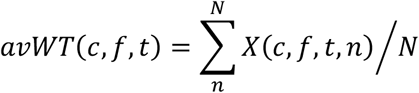

where *X*(*c, f, t, n*) is the time-frequency coefficient at channel *c*, frequency *f*, time *t* and segment *n* of the EEG signal given by *X*(*c, t, n*). Following the ERPWAVELAB data structure (Mørup et al., 2007), the resulting MATLAB variable “WT” is a 3D matrix (Channel × Frequency × Time) that contains the average of the transformed segments. Notice that, since WTools takes the modulus of the complex coefficients, the data are still expressed in *μ*V rather than in power (i.e. *μ*V^2^). As a result, the measure is slightly more sensitive to small frequency changes.

WTools allows computing both total-induced and evoked oscillations (David et al., 2006; Tallon-Baudry et al., 1996). Total-induced oscillations are computed by wavelet transformation of each epoch before averaging all of them together (i.e. the pipeline we just described above): in this way, the oscillatory activity non-phase-locked to the onset of the stimulus is preserved. Total-induced oscillations are richer measures compared to evoked oscillations. Evoked oscillations are computed by running the wavelet transformation after averaging all the epochs together, that is the wavelet transformation is performed on the average (i.e. it is the wavelet transformation of the ERP): in this way, the oscillatory activity that is non-phase-locked to the stimulus onset is averaged out and lost before the wavelet transformation. Evoked oscillations are poorer measures compared to total-induced oscillations and are slightly more sensitive to low-level features of the stimulus.

#### 2.1.3 Edges chopping

The continuous wavelet transformation assumes signal continuity, yet EEG segments inherently lack this continuity: at the segment’s start, a shift from no signal to signal with amplitude occurs, mirrored at the segment’s end, causing significant frequency changes. This discontinuity distorts coefficient vectors at the edges, particularly noticeable at lower frequencies where wavelets are broader (see Fig. 2). To address the impact of this distortion on EEG signal analysis, it is advisable to pre-cut wider segments. Alternatively, WTools offers an edge padding feature to compensate for signal absence at segment edges. Based on empirical evaluation, the minimally necessary length of extra signal needed at each edge can be estimated with the current formula (Gergely Csibra, personal communication):

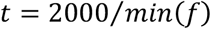

where *f* is the frequency range of interest. For instance, if one wants to look at frequencies between 5-15 Hz, then it is necessary to consider *t* = 400 ms of extra signal at both left and right edges of the segment.

### 2.1.4 Baseline correction

Differently from other toolboxes that adopt a divisive baseline (Grandchamp and Delorme, 2011), WTools performs a subtractive baseline similarly to baseline correction for ERPs. The average amplitude in a given time window of the pre-stimulus interval is computed and subtracted from the whole time-varying signal. Notice that there is not a common baseline for all frequencies; instead, each frequency of interest has its own baseline value that is used for correction only for that frequency. This approach makes the resulting spectrograms easy to read and highly comparable to ERPs, which is particularly handy for developmental research. Furthermore, adopting a divisive baseline assumes a multiplicative model, in which oscillations and 1/*f* activity scale by the same factor relative to baseline for each frequency (Gyurkovics et al., 2021). This assumption may be incorrect when signal and noise contribute independently to the power spectrum (i.e. additive model; Gyurkovics et al., 2021), a very common problem in developmental studies for which a subtractive baseline may be more suitable.

### 2.2 Dataset description

To examine the time-frequency functionality of WTools, we employed a dataset from a previously published study investigating EEG responses to multimodal ostensive signals in 5-month-old infants (Experiment 1 in Parise and Csibra, 2013). The study was approved by the United Ethical Review Committee for Research in Psychology (EPKEB) at Central European University, and the parents of all participants provided written informed consent. Eighteen infants (9 females; average age= 148.17 days, range = 136 to 157 days) were included in the study. All infants were born full-term (gestational age: 37 to 41 weeks) and in the normal weight range (>2500 g). We provide free access to an anonymized dataset containing three representative participants from this previously published study (see section “WTools availability”). The figures contained in the present paper pertain to one of these participants.

The experiment conformed to a 2×2 within-subject factorial design, corresponding to the orthogonal crossing of the factors Modality (visual vs. auditory) and Ostension (ostensive vs. non-ostensive). As a result, the experiment included four within-subject conditions: ostensive visual stimulus of direct gaze (DG); non-ostensive visual stimulus of averted gaze (AG); ostensive auditory stimulus of infant-directed speech (IDS); non-ostensive auditory stimulus of adult-directed speech (ADS).

Visual stimuli (female faces) were presented on a 19-inch CRT monitor operating at 100 Hz refresh rate using PsychToolBox (v. 3.0.8) and custom-made MATLAB scripts. Auditory stimuli (pseudo-words pronounced by a female voice) were presented by a pair of computer speakers located behind the monitor. A remote-controlled video camera located below the monitor allowed the recording of infants’ behaviour during the experiment. For further details on the stimuli and setup, please refer to Parise and Csibra, 2013.

Infants sat on their parent’s lap at 70 cm from the monitor. At the beginning of each trial, a dynamic visual stimulus appeared on top of the face with closed eyes for 600 ms to grab the infant’s attention. Afterwards, the attention grabber stopped moving and stayed on screen for a random interval between 600 and 800 ms. The attention grabber then disappeared and a visual (DG or AG) or auditory (IDS or ADS) stimulus was presented for 1000 ms. An inter-trial interval between 1100 and 1300 ms was inserted between successive trials. Infants were presented with a maximum of 192 trials divided into 4 blocks. The behaviour of the infants was video-recorded throughout the session for offline trial-by-trial editing.

High-density EEG was recorded continuously using Hydrocel Geodesic Sensor Nets (Electrical Geodesics Inc., Eugene, OR, USA) at 124 scalp locations referenced to the vertex (Cz). The ground electrode was at the rear of the head (between Cz and Pz). Electrophysiological signals were acquired at the sampling rate of 500 Hz by an Electrical Geodesics Inc. amplifier with a band-pass filter of 0.1–200 Hz.

### 2.3 Data analysis

The digitized EEG was band-pass filtered between 0.3-100 Hz and was segmented into epochs including 500 ms before stimulus onset and 1500 ms following stimulus onset for each trial. EEG epochs were automatically rejected for body and eye movements whenever the average amplitude of an 80 ms gliding window exceeded 55 µV at horizontal EOG channels or 200 µV at any other channel. Additional rejection of bad recording was performed by visual inspection of each individual epoch. Bad channels were interpolated in epochs in which ≤10% of the channels contained artefacts; epochs in which >10% of the channels contained artefacts were rejected. Infants contributed on average 12.11 artefact-free trials to the DG condition (range: 10 to 19), 11.67 to the AG condition (10 to 15), 11.67 to the IDS condition (10 to 19), 12.61 to the ADS condition (10 to 22). The artefact-free segments were subjected to time-frequency analysis using WTools and EEGLAB respectively.

#### 2.3.1 WTools analysis

To import data into WTools, epochs were transformed from the native EGI format to the MATLAB format using the appropriate NetStation Waveform export tool by EGI, and then the correspondent EEGLAB import plugin (v. 2021.1). In general, WTools calls existing EEGLAB plugins to import data. Each plugin takes as input a specific data format. Users must save the data in a format that is compatible with the EEGLAB plugin of interest, which is called by the WTools import routine (see guidelines: eeglab.org/tutorials/04_Import/Import.html). Note that data to be imported into WTools must conform to an event-related within-subject design with one single trigger per segment/epoch indicating the onset of the key event. Epochs were re-referenced to average reference and then subjected to time-frequency analysis. Data were analysed using both the standard WTools pipeline (whose code is released with this publication) and custom pipelines that allowed a direct comparison with the EEGLAB output, as outlined below.

In the following, we describe the standard WTools pipeline. We computed complex Morlet wavelets for the frequencies 10-90 Hz in 1 Hz steps. We calculated total-induced oscillations by performing a continuous wavelet transformation of all the epochs using convolution with each normalized wavelet (number of cycles = 7) and taking the absolute value (i.e., the amplitude) of the results (Csibra et al., 2000). To remove the distortion introduced by the convolution, we chopped 300 ms at each edge of the epochs, resulting in 1400 ms long segments, including 200 ms before and 1200 ms after stimulus onset. We used the average amplitude of the 200 ms pre-stimulus window as baseline, subtracting it from the whole epoch at each frequency.

To compare the WTools output to the EEGLAB output, we carried out a custom analysis that deviated from the standard WTools pipeline in two ways. First, we took the squared value (i.e. the power) of the complex wavelet coefficient, in line with the EEGLAB pipeline. Second, we skipped baseline correction to avoid a mismatch between the two toolboxes (while WTools performs a subtractive baseline, EEGLAB implements a divisive baseline).

### 2.3.2 EEGLAB analysis

Epochs were transformed from the native EGI format to the MATLAB format using the NetStation export tool and then imported by using the correspondent EEGLAB import plugin (v. 2021.1). Epochs were re-referenced to average reference and then subjected to time-frequency analysis. Data were analysed using various EEGLAB pipelines that allowed a direct comparison with the WTools output. For all analyses, we computed Morlet complex wavelets for the frequencies 10-90 Hz in 1 Hz steps. We calculated total-induced oscillations by performing a continuous wavelet transformation of all the epochs using convolution with each normalized wavelet (number of cycles = 7) and taking the squared value (i.e., the power) of the complex coefficients. Data were automatically chopped by EEGLAB to remove the distortion introduced by the convolution, resulting in 1400 ms long segments, including 200 ms before and 1200 ms after stimulus onset. For baseline correction, we entered the 200 ms pre-stimulus window as baseline.

To compare the EEGLAB output to the WTools output, we carried out a custom analysis that deviated from the standard EEGLAB pipeline in two ways. First, we did not log-transform the results of the wavelet transformation (typically done to bring the results to the standard dB scale). Second, we skipped baseline correction to avoid a mismatch between the two toolboxes.

## 3 Results

### 3.1 Illustrating WTools

We illustrate the main visualization functionalities available in WTools using results obtained under the standard WTools pipeline. For details of all the available visualization settings, please refer to the step-by-step tutorial of the entire pipeline that is provided on GitHub (see section “WTools availability”). Three main types of plots are available: time-frequency graphs, 2D scalp maps and 3D scalp maps. For time-frequency graphs, it is possible to plot results from individual channels, from the average of a desired subset of channels, or from all available channels in a topographical arrangement. For each of these graphs, users can also plot the standard error of the effect relative to baseline, which allows a preliminary descriptive evaluation of the effect size. For 2D and 3D scalp maps, please note that WTools expects to assign a 2D topoplot configuration to any given dataset during the initial data import routine. Optionally, the user can decide to additionally select a spline configuration for 3D scalp maps. Currently, WTools offers several 2D and 3D configuration files (ANT, BIOSEMI, GSN) for 2D and 3D maps of adult and infant data. It is possible to integrate and extend the toolbox with personalized configurations for developmental research (i.e. newborn and infant scalp maps). For 2D scalp maps, the user can obtain a plot corresponding to various possibilities: one timepoint at one specific frequency; average of timepoints at a specific frequency; average of a frequency range at one timepoint. It is also possible to plot a series of scalp maps, either for time or frequency series. For 3D scalp maps, the user can similarly obtain a plot corresponding to various possibilities: one timepoint at one certain frequency; average of timepoints at a certain frequency; average of a frequency range at one timepoint. Finally, all types of graphs allow plotting results of individual conditions or results of user-defined pair-wise differences between conditions.

For illustrative purposes, we show the time-frequency plots and 2D scalp maps for one representative participant and channel both for individual conditions and for their difference (Fig. 3). Furthermore, we show the 3D infant scalp map of the same difference (Fig. 4). Results show a higher synchronization for Condition 1 (DG) than Condition 2 (AG) in the gamma frequency-band at 250-350 ms post-stimulus. This result is representative of the final finding of the original paper (Parise and Csibra, 2013), which replicates and extends previous work (Grossmann et al., 2007). Time, frequency and scale ranges are user-defined before all plotting.

**Fig. 3.**
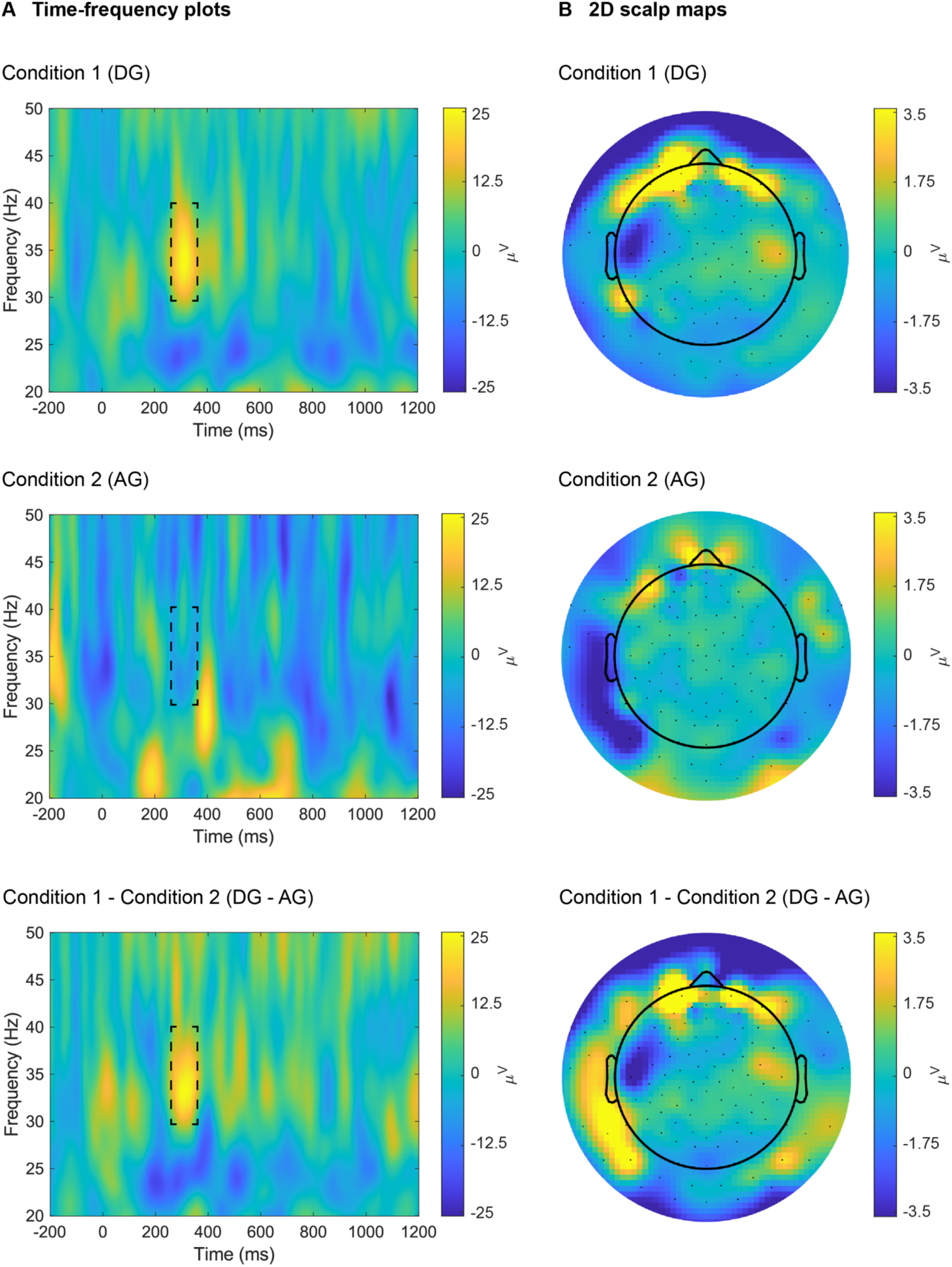
Time-frequency plots (**A**) and 2D scalp maps (**B**) for one representative participant (“04”) and channel (“22”, corresponding to Fp2) for individual conditions (“DG”, “AG”) and for their difference (“DG – AG”), under the standard WTools pipeline. Scalp maps show the topographies of the time-frequency results in the 250-350 ms time window and 30-40 Hz frequency window (as highlighted with dashed lines in panel A).

**Fig. 4.**
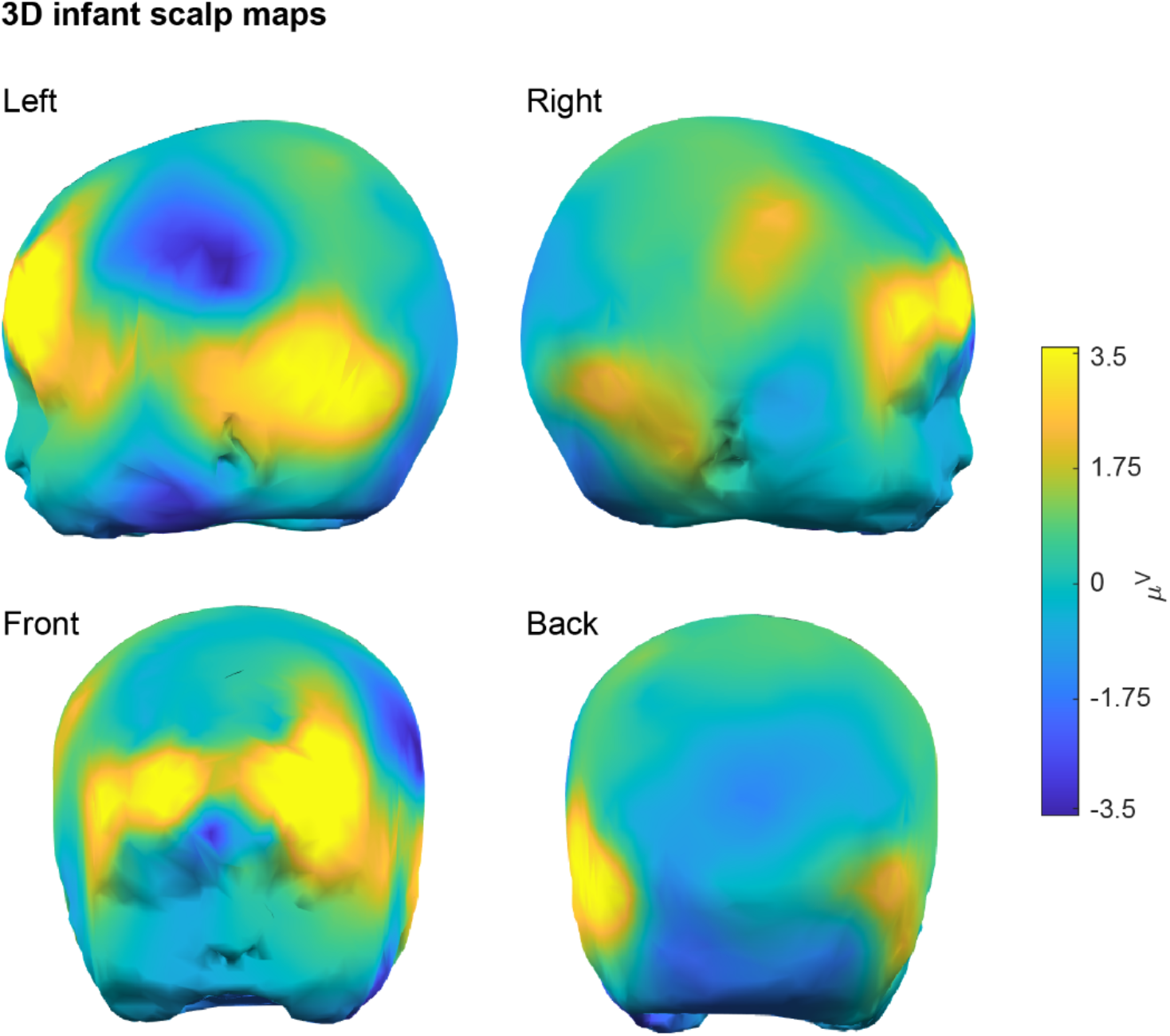
3D infant scalp maps for the pair-wise condition difference “DG – AG” of one representative participant (“04”) under the standard WTools pipeline. Scalp maps show the topographies of the time-frequency results in the 250-350 ms time window and 30-40 Hz frequency window (as highlighted in Fig. 3A). In the GUI, users can rotate scalp maps in any directions to optimize scalp visualization as desired.

Finally, WTools has a built-in functionality that allows exporting summary data for later statistical inference. Specifically, users can extract numeric values for each desired subject, condition and channel, computing the average value both in a time window and in a frequency range (via a double average). Values are stored in a text-tabulated file (.tab) that can be treated directly by third-party statistical packages.

### 2.3 Comparison with EEGLAB

The analysis aimed to examine the time-frequency functionality of WTools via comparison with state-of-the-art methods implemented in EEGLAB, an established popular free software (Delorme and Makeig, 2004). First, we report the time-frequency results for one representative participant (“04”), channel (“22”, corresponding to Fp2) and condition (“DG”) under the custom pipelines that allowed a direct comparison of the WTools and EEGLAB outputs. Second, we report the time-frequency results under the standard WTools and EEGLAB pipelines, which are practically used in real research settings.

By removing baseline correction and taking the squared value of the complex wavelet coefficient (see section “Data analysis” for further details), the WTools output perfectly matched the EEGLAB output (Fig. 5A). This analysis confirms that WTools produces time-frequency results in line with state-of-the-art EEGLAB outputs (our gold standard for the present work), representing an important sanity check. Then, as expected, the standard pipelines generated slightly different results across WTools and EEGLAB (Fig. 5B). Importantly, the two toolboxes still showed qualitatively compatible results, with a pronounced increase of induced gamma-band activity over the forehead (channel 22, corresponding to Fp2) in the 250 to 350 ms time window, and 30 to 40 Hz frequency window (Parise and Csibra, 2013). Please see the Supplementary material for a comprehensive review of the results under additional analyses pipelines, which derived from the orthogonal crossing of the settings being manipulated (i.e. baseline correction, complex wavelet manipulation, log-transformation).

**Fig. 5.**
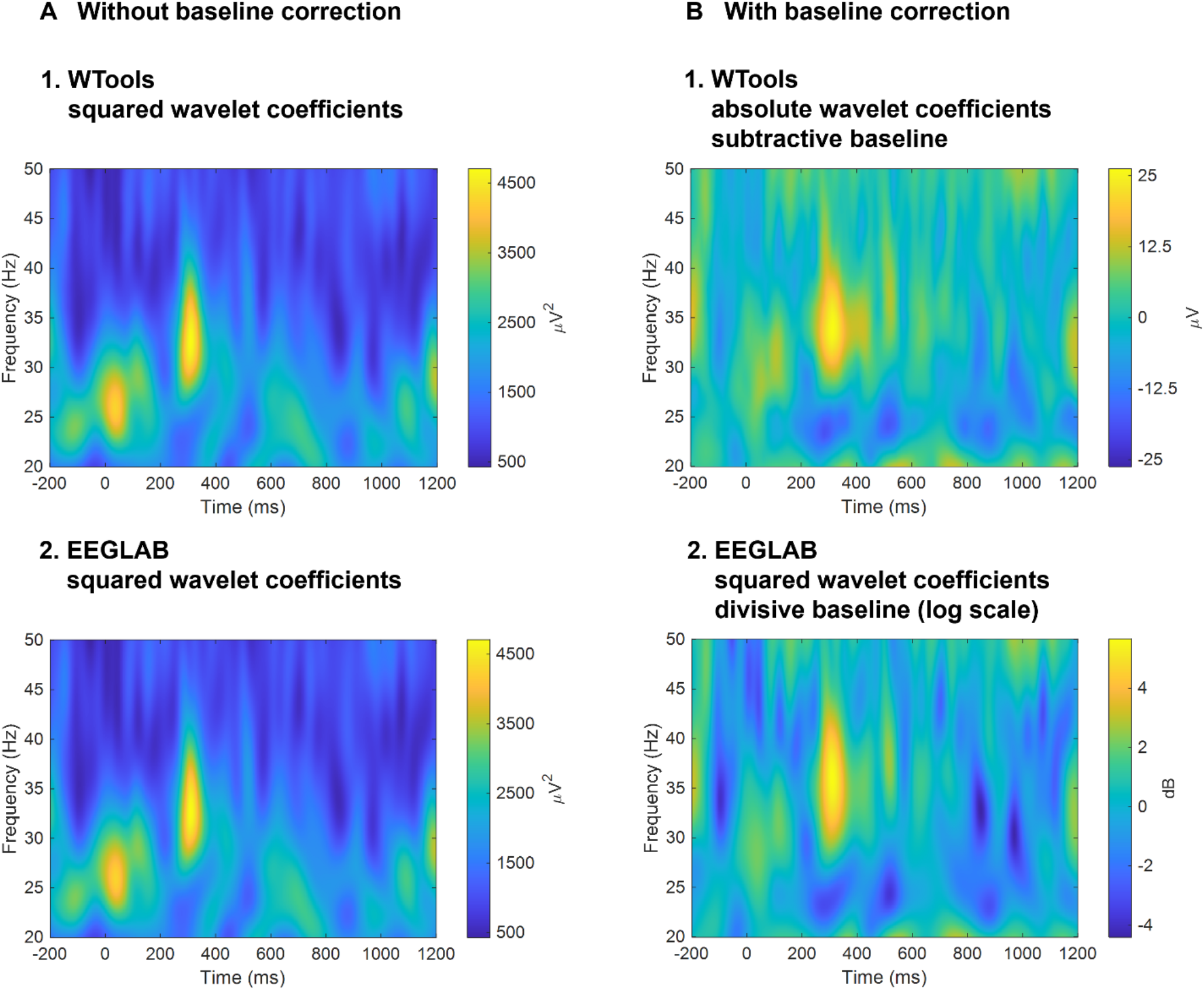
Time-frequency plots for one representative participant (“04”), channel (“22”, corresponding to Fp2) and condition (“DG”) under the custom pipelines that allowed a direct comparison of the WTools and EEGLAB outputs (**A**) and under the standard WTools and EEGLAB pipelines (**B**).

## 4 Discussion

This paper introduced WTools, a new MATLAB-based toolbox for time-frequency analysis of the EEG signal that implements complex wavelet transformation. We provided a detailed description of the WTools algorithm for wavelet transformation, as well as an overview of the main visualization functionalities for data exploration and characterization. Crucially, we compared WTools with state-of-the-art methods implemented in EEGLAB (Delorme and Makeig, 2004). Finally, we created a step-by-step illustrated and user-friendly tutorial of the entire processing pipeline for easy and transparent access to the toolbox’s functionalities (see section “WTools availability”).

In terms of usability, WTools features a straightforward, transparent and user-friendly GUI that is particularly handy for non-advanced users (with little to no experience in coding). Furthermore, WTools guarantees high flexibility in data handling, since it takes a huge range of input formats thanks to its direct compatibility with EEGLAB: any dataset that can be imported into EEGLAB can also be imported into WTools.

In terms of data analysis, WTools produces time-frequency results in units of *μ*V and it performs a subtractive baseline correction, very similar to baseline correction for ERPs. This approach makes the resulting spectrograms easy to read and highly comparable to ERPs, which is particularly handy for developmental research. Plotting functions allow for a straightforward visualization and evaluation of the data in terms of time-frequency graphs, 2D and 3D scalp maps (including both infant and adult 3D configurations). Users can also plot the standard error of an effect relative to baseline, which allows a preliminary descriptive evaluation of the effect size. Finally, WTools features a built-in tool for exporting numerical data that can be treated directly by third-party statistical packages.

For these reasons combined, an increasing number of studies in the field of developmental cognitive neuroscience have adopted beta versions of WTools (de Klerk et al., 2016; Kolesnik et al., 2019; Ortiz-Barajas et al., 2023; Piccardi et al., 2020; Pomiechowska and Csibra, 2017; Quadrelli et al., 2021, 2019; Southgate and Vernetti, 2014; Yin et al., 2020). With the current official release, we further foster the transparent and reproducible usage of WTools by the developmental cognitive neuroscience community and beyond.

## WTools availability

WTools is publicly available as open-source software on GitHub (github.com/cogdevtools/WTools/tree/v2.0) under GNU General Public License (GPL-3.0). A step-by-step tutorial for the use of WTools is provided in the wiki section of the GitHub project. An example anonymized dataset (data from Parise and Csibra, 2013) has been deposited in the Open Science Framework and is freely available at osf.io/jtudr.

## Acknowledgements

When developing WTools, E.P. and L.F. were supported by an Advanced Investigator Grant (#249519, OSTREFCOM) from the European Research Council to Gergely Csibra. E.P. was partially supported by the International Centre for Language and Communicative Development (LuCiD) at Lancaster University, funded by the Economic and Social Research Council (UK) [ES/L008955/1]. We thank John E. Richards for providing E.P. with the MRI scan to create the 3D infant head model embedded in WTools. We thank Teodora Gliga and Morgan Whitworth for testing preliminary versions of this WTools release.

