## Supplementary Figure S1 for "WTools: a MATLAB-based toolbox for time-frequency analysis"

### Supplementary materials

Ambra Ferrari[a*], Luca Filippin[b], Marco Buiatti[c], Eugenio Parise[d*]

[a] CIMeC, Center for Mind/Brain Sciences, University of Trento, Corso Bettini 31, 38068, Rovereto, Italy;
Max Plank Institute for Psycholinguistics, Wundtlaan 1, 6525 XD, Nijmegen, The Netherlands;


[b] CIMeC, Center for Mind/Brain Sciences, University of Trento, Corso Bettini 31, 38068, Rovereto, Italy;


[c] CIMeC, Center for Mind/Brain Sciences, University of Trento, Piazza Manifattura 1, 38068, Rovereto, Italy;


[d] CIMeC, Center for Mind/Brain Sciences, University of Trento, Corso Bettini 31, 38068, Rovereto, Italy;


[*]Corresponding authors

**Fig. S1** | Comprehensive overview of the time-frequency results under all analyses pipelines, which derived from the orthogonal crossing of the manipulated settings (i.e. baseline correction, complex wavelet manipulation, log-transformation). We ran four time-frequency analyses in WTools corresponding to the orthogonal crossing of 1) baseline correction (with or without subtractive baseline) and 2) complex coefficient transformation (absolute or squared value). We ran four time-frequency analyses in EEGLAB corresponding to the orthogonal crossing of 1) baseline correction (with or without divisive baseline) and 2) log-transformation of the time-frequency results to convert them to the standard dB scale (with or without log-transformation). Time-frequency plots correspond to one representative participant ("04"), channel ("22", corresponding to Fp2) and condition ("DG").

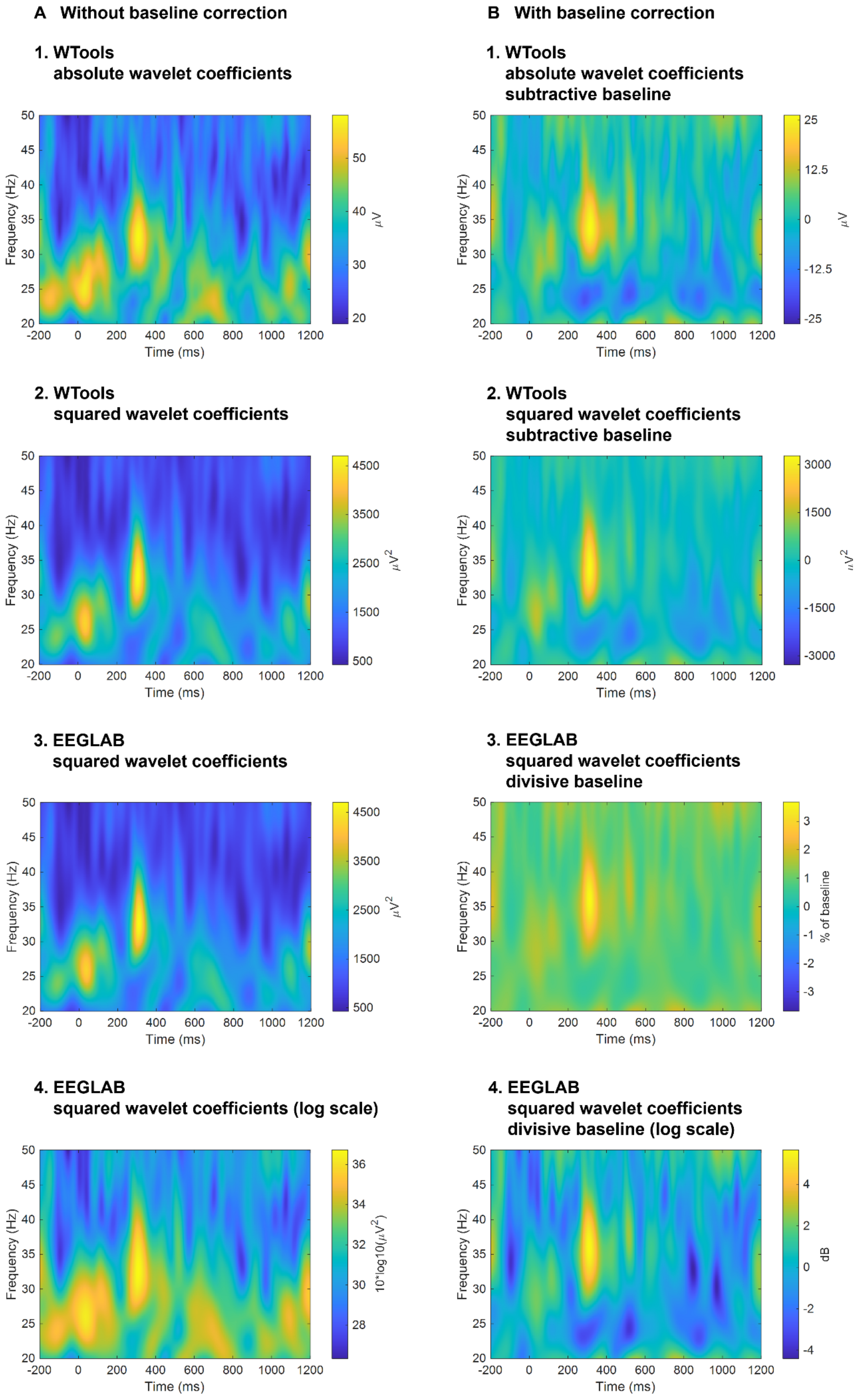
A Without baseline correction
B With baseline correction
1. WTools
absolute wavelet coefficients
1. WTools
absolute wavelet coefficients
subtractive baseline
2. WTools
squared wavelet coefficients
2. WTools
squared wavelet coefficients
subtractive baseline
3. EEGLAB
squared wavelet coefficients
3. EEGLAB
squared wavelet coefficients
divisive baseline
4. EEGLAB
squared wavelet coefficients (log scale)
4. EEGLAB
squared wavelet coefficients
divisive baseline (log scale)
Frequency (Hz)
Time (ms)
μV
μV2
% of baseline
10*log10(μV2)
dB
